# Development of molecular markers for western honey bee (*Apis mellifera* L.) subspecies of regulatory concern in the United States

**DOI:** 10.64898/2026.03.05.709340

**Authors:** Jose A. P. Marcelino, Leigh Boardman, Matthew R. Moore, James C. Fulton, Hector Urbina, Laura Patterson Rosa, Bernd Grünewald, Mike Allsopp, Christian Pirk, Garth Cambray, James D. Ellis

## Abstract

Rapid, accurate and cost-effective identification of *Apis mellifera* subspecies is needed for subspecies of regulatory concern. We designed and validated subspecies markers based on single nucleotide polymorphisms (SNP) on mitochondrial *cytochrome b* (*Cytb*) and *NADH dehydrogenase 4* (*ND4*) genes. We used a combination of established and novel real-time qPCRs in a stepwise series, with increasing discrimination power, to (1) differentiate honey bees of African (A-lineage) ancestry from those of other lineages (*Cytb* SNP #1), (2) identify African-derived honey bees (AHBs) (*Cytb* SNP #2), and (3) detect *A. m. capensis* exclusively in the bee’s indigenous region of South Africa (*ND4* SNP). We also developed a restriction fragment length polymorphism assay targeting a SNP on the *NADH dehydrogenase 2* (*ND2* RFLP) gene to detect the specific mitochondrial A-lineage clade. These assays allow for reliable time- and cost-effective results that provide increased accuracy on subspecies assignation.

## 1. Introduction

The ability to identify subspecies of *Apis mellifera* Linnaeus, the western honey bee, accurately, timely, and cost effectively is important for research and regulatory purposes. From a regulatory perspective, some honey bee subspecies that have moved outside their native ranges are considered problematic in their introduced ranges, heightening the need to have accurate diagnostic measures for these subspecies. Developing a reliable method to identify honey bee subspecies has been difficult due to the ongoing movement of honey bee stocks globally, largely for beekeeping purposes, which has resulted in significant genetic admixture within the subspecies (e.g., Harpur et al. 2012). This is exacerbated in places like the United States of America (USA) where 10+ honey bee subspecies or stocks have been introduced (Seeley 2019; Carpenter and Harpur 2021; Marcelino et al. 2022).

The USA falls within the invasive range of *A. m. scutellata* Lepeletier, a subspecies that has had a tremendous economic and societal impact throughout the Americas and one that can be difficult to identify accurately (Marcelino et al. 2022). Stocks of honey bees derived from *A. m. scutellata*, a subspecies native to southern and eastern Africa, originated in Brazil from the escape of *A. m. scutellata* from a research breeding program. After its release, the bee hybridized with local *A. m. ligustica* Spinola and *A. m. mellifera* Linnaeus of European ancestry already in Brazil (Kerr 1992). The resulting hybrid, originally called the Africanized honey bee (Gonçalves 1974; Kerr 1992), spread widely across the Americas. It was found in the USA in 1990 (Caron 2001) and now occurs in several states (Carpenter and Harpur 2021). Generally, African-derived honey bees (AHBs) pose a greater risk to humans through higher risk of sting-related incidents and heightened defensive behavior than do the European-derived honey bee stocks (EHBs) managed by beekeepers in the USA. AHBs also pose a risk to managed honey bee colonies as they can outcompete EHBs where they co-occur (reviewed in Breed et al. 2004; Schneider et al. 2004). Correspondingly, many states in the USA have regulations against the management of AHBs (Marcelino et al. 2022).

*Apis mellifera capensis* Eschscholtz is another subspecies of regulatory concern in the USA. Naturally occurring in the Cape region of South Africa, this subspecies frequently exhibits thelytokous parthenogenesis (Onions 1914; Goudie and Oldroyd 2014; Mumoki et al. 2021), or the ability of worker honey bees to lay unfertilized diploid eggs that develop into adult female bees (workers and queens). A clonal subpopulation of these workers is causing damage to the commercial hives of *A. m. scutellata* in South Africa (Pirk et al. 2014). While the subspecies is not currently present in the USA, the risk of accidental introduction led to the subspecies being listed in the U.S. Code of Federal Regulations 7 CFR Parts §322.1 (US Code of Federal Regulation, 7-B-III-322.).

At present, multiple techniques exist for honey bee subspecies assignment. Legacy methods for identification include geomorphometric or morphometric measurements of wing vein positions (Rinderer et al. 1986), which were quickly complemented with the development of molecular methods (reviewed in Meixner et al. 2013). Newer diagnostic techniques tend to use whole-genome single nucleotide polymorphisms (SNP) approaches (e.g., Whitfield et al. 2006; Avalos et al. 2017; Donthu et al. 2024, and references therein). However, these methods are inaccessible to some researchers due to high cost and time investment, and undesirable in circumstances when a prompt identification is required for regulatory purposes, e.g. intercepts at ports of entry or apiary queen replacements. Assays may also be limited by a lack of reference material for other African subspecies that are similar to *A. m. scutellata*, and thus more likely to result in false positives. Furthermore, genomic data do not always result in clearer discrimination, especially when trying to identify a subspecies characterized by a behavioral phenotype – see discussions on thelytoky in *A. m. capensis* (Aumer et al. 2019; Christmas et al. 2019; Yagound et al. 2020). Members of our author team have specifically investigated samples of *A. m. capensis*, *A. m. scutellata*, and hybrids from South Africa using a variety of these methods: morphometrics, microsatellites, mitochondrial genomes, and SNPs (Eimanifar et al. 2018a, 2018b, 2020; Bustamante et al. 2020; Patterson Rosa et al. 2021).

Our goal is to contribute to existing efforts aimed at developing comprehensive, accurate and time-effective genetic tests for AHBs and *A. m. capensis.* Recently, we developed a quantitative PCR (qPCR) assay targeting a SNP in *cytochrome b* gene (*Cytb*) that can differentiate honey bees from the African A-lineage (Boardman et al. 2021) from those of other lineages based on pre-existing restriction fragment length polymorphisms assays (Crozier et al. 1991; Pinto et al. 2003; Szalanski and McKern 2007) and mitochondrial sequence data (Pinto et al. 2007). This assay can identify maternally African-derived AHBs, but it cannot discern among nine different African subspecies in the African (A-) clade. Here, we demonstrate a more specific AHB qPCR assay – also based on a *Cytb* SNP. Furthermore, we developed a *NADH-dehydrogenase subunit 4* (*ND4*) gene qPCR assay to detect *A. m. capensis* based partially on data from Eimanifar et al. (2018b). Lastly, to provide an alternative to qPCR technology, we developed a *NADH-dehydrogenase subunit 2* (*ND2*) RFLP assay to detect bees with maternal *A. m. adansonii* Latreille*, A. m. capensis, A. m. monticola* Smith, and *A. m. scutellata* ancestry (referred to as the ACMS clade to correspond with *adansonii*, *capensis*, *monticola*, *scutellata*) based on the discriminatory power of this gene for *A. mellifera* subspecies (Arias and Sheppard 1996; Ilyasov et al. 2011).

## 2. Material and methods

### 2.1. Background information and pilot data

Mitochondrial genes were chosen to screen for SNPs of interest based on the ability of previous data to resolve *A. mellifera* subspecies. The target genes included *Cytb* (Crozier et al. 1991; Pinto et al. 2003; Szalanski and McKern 2007; Boardman et al. 2021), *ND2* (Arias and Sheppard 1996; Ilyasov et al. 2011), and *ND4* (Eimanifar et al. 2018b).

All available *A. mellifera Cytb* (n=226) and *ND2* (n=241) sequences were downloaded from NCBI GenBank (ncbi.nlm.nih.gov) and merged with a dataset of 91 mitochondrial genomes (Boardman et al., in preparation). This resulted in a dataset that comprised 317 *A. mellifera Cytb* sequences and 332 *A. mellifera ND2* sequences (Electronic Supplementary Materials Data S1 and Data S2). *Apis cerana* sequences from mitochondrial genome GQ162109 (Tan et al. 2011) were used as outgroups for further analyses. These alignments were used to generate phylogenetic trees, and the analyses recovered clades that directed the further development of the assays described herein (Figure S1, Figure S2). We used the author-assigned subspecies from the original published work in these alignments and phylogenies.

Ongoing use of the A-lineage assay (referred to hereafter as *qPCR assay I*, Boardman et al., 2021) by the Florida Department of Agriculture and Consumer Services - Division of Plant Industry (FDACS - DPI) identified several defensive honey bee colonies in Florida, submitted from stinging reports and other behavioral complaints, that were positive for the A-lineage marker. Follow-up Sanger sequencing of *Cytb* was performed on these A-lineage samples. This sequencing effort revealed a SNP at position 146 (based on the *A. m. ligustica* positions used in Crozier and Crozier (1993) and Pinto et al. (2007)), that was also found in the mitochondrial genome reference honey bee sequences from Mexico (KJ601784, Gibson and Hunt 2016), Brazilian bees (EF184030.1, Pinto et al. 2007), AHB haplotype 3 from Arkansas, USA (EF016648, Szalanski and McKern 2007), Mbo-A/Hinf-1(1) haplotype (EU513284, Ferreira et al. 2009), and some *A. m. scutellata* from locations in South Africa (MT572364-MT57239; MW530563-MW530565, see Boardman et al. 2021). This candidate SNP was then targeted for the development of *qPCR assay II* to detect AHBs.

While samples from South Africa can be difficult to discern since *A. m. scutellata* and *A. m. capensis* co-exist in an overlapping hybrid zone (c.f. Hepburn et al. 1998; Eimanifar et al. 2018a, 2020; Patterson Rosa et al. 2021), several mitochondrial genomes from the native Cape region of *A. m capensis* in South Africa form a distinct clade that could be considered the most representative of *A. m. capensis* (Eimanifar et al. 2018b). We developed a qPCR marker for the mitochondrial *ND4* gene region that can discriminate *A.m. capensis* bees from this region from all other subspecies.

### 2.2. Assay development and design overview

Our series of qPCRs (Figure 1) aimed to differentiate honey bees of mitochondrial African ancestry, AHBs, and *A. m. capensis*. This was designed to increase specificity and resolution power of assignations by relying on a comprehensive sample dataset (Table 1) that would help to eliminate false positives caused by north African and Mediterranean A-lineage subspecies. The assay series comprises (1) expanded sample testing of the *qPCR assay I* (Boardman et al. 2021); (2) a new *qPCR assay II* to detect AHBs; and (3) a new *qPCR assay III* to detect *A. m. capensis.* Real-time *qPCRs assays II* and *III* were validated by Sanger sequencing of *Cytb* and *ND4*. Lastly, we developed a low cost RFLP marker targeting a restriction enzyme site in the mitochondrial *ND2* gene that discerns honey bees with maternal *A. m. adansonii, A. m. capensis, A. m. monticola*, *A. m. scutellata* (referred to collectively as ACMS clade) ancestry from all other subspecies. This can be used when qPCR technologies are not available or as a secondary assay confirming qPCR results.

**Figure 1.**
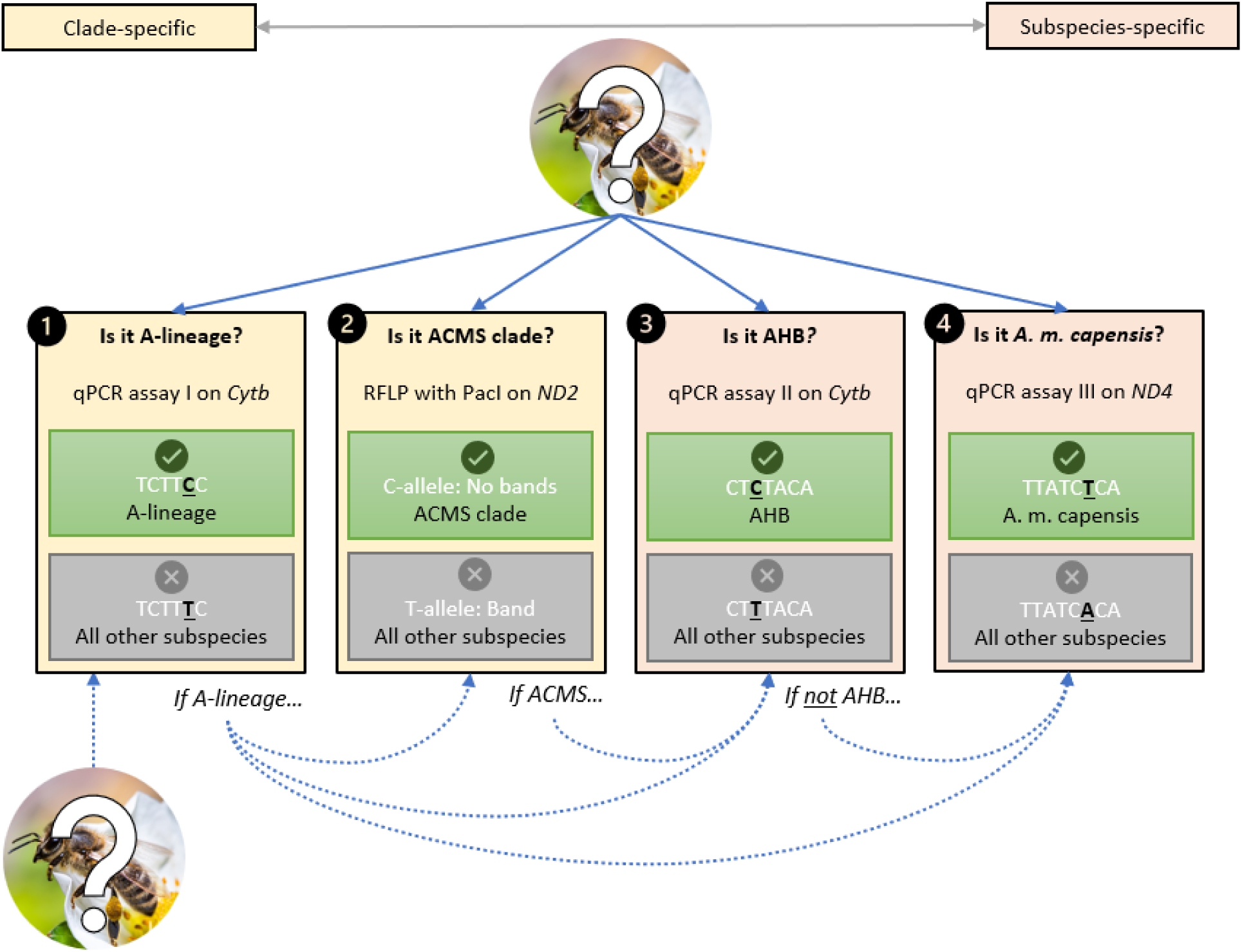
Protocols to establish the maternal ancestry of an unknown *Apis mellifera* honey bee using clade-specific and/or subspecies-specific assays based on a single SNP. The qPCR assays targeting single SNPs (bold, underlined) to identify: African A-lineage (1), possible African-derived honey bees (AHBs) (3), possible *A. m. capensis* (4). An RFLP assay (2) for differentiating *A. m. adansonii, A. m. capensis, A. m. monticola*, *A. m. scutellata* ancestry (referred to as ACMS clade) provides an alternative for labs without access to qPCR technology. All results were validated with Sanger sequencing. Assays can be used independently (solid arrows) or as a pipeline protocol (dashed arrows).

**Table 1.** Summary of sampling information. Honey bee (*Apis mellifera*) samples (n=145) were obtained from the Ruttner Bee Collection at the Bee Research Institute in Oberursel, Germany (numerical sample ID used by Institute staff) or collected from South Africa or USA (Arizona, Florida, Puerto Rico and Texas) (two letter sample ID). Please refer to Table S1 for full details on these samples.

| n | Sample ID | Note | Country | Location |
| --- | --- | --- | --- | --- |
| Genomic DNA leftover from (Boardman et al. 2021) |  |  |  |  |
| 15 | 88-102 | Specimens from fourteen different *A. mellifera* subspecies (Oberursel collection) | Various | Various |
| 18 | 67-84 | *A. m. capensis* and *A. m. scutellata* (HBREL collection) | South Africa | Various |
| 3 | 85-87 | Immatures (queen) | USA | Florida |
| Individual honey bees |  |  |  |  |
| 35 | 1-29, 103-108 | *A. m. capensis*, *A. m. scutellata* and hybrid specimens (HBREL collection) | South Africa | Various |
| 37 | 30-66 | Individual bees collected from colonies responsible for stinging incidents (FDACS) | USA | South Florida |
| 16 | 113-115, 129-132, 137-145 | Defensive honey bees. Suspected AHB | USA | Arizona |
| 6 | 109, 110, 125, 126, 133, 134 | Defensive honey bees. Suspected AHB | USA | Texas |
| 3 | 116-118 | Gentle honey bees | USA | Florida |
| 6 | 111, 112, 127, 128, 135, 136 | Gentle African and European ancestry specimens, and defensive honey bees | USA | Puerto Rico |
| 3 | 116-118 | Gentle honey bees | USA | Florida |
| 1 | 119 | Feral, defensive honey bees. Suspected AHB | USA | North Florida |
| 4 | 120-123 | FABIS identified samples | USA | Florida |
| 1 | 124 | Positive control for honey bee of European ancestry. Pooled sample of three bees | USA | Florida |

n = number of individual bees sampled; AHB = African-derived honey bee; HBREL = University of Florida Honey Bee Research and Extension Laboratory; FDACS = Florida Department of Agriculture and Consumer Services.

While each qPCR assay can be used independently, they were designed to be used as a series of qPCRs that offer increasing specificity (Figure 1). Results are also more robust if multiple assays are used on the same sample. Detailed information about each assay follows.

#### 2.2.1. New qPCR assay II – AHB

*qPCR assay II* amplifies a region of *Cytb* and fluorescently detects an internal SNP discriminating AHBs from all others honey bees (Figure 1) using a dual-labeled Locked Nucleic Acid (LNA) probe. Primers were placed in regions of highly conserved sequence homology, with similar annealing temperatures (≤ 2 °C difference), and as close to the target SNP as possible. Primers and probes (Table 2) were tested for off-target matches using Basic Local Alignment Search Tool (BLAST) of the oligonucleotides against *Apis* sequences in the NCBI database. A Ct value (fluorescence) indicates a C-allele, and possible AHB. Undetermined Ct values indicate a T-allele.

**Table 2.**
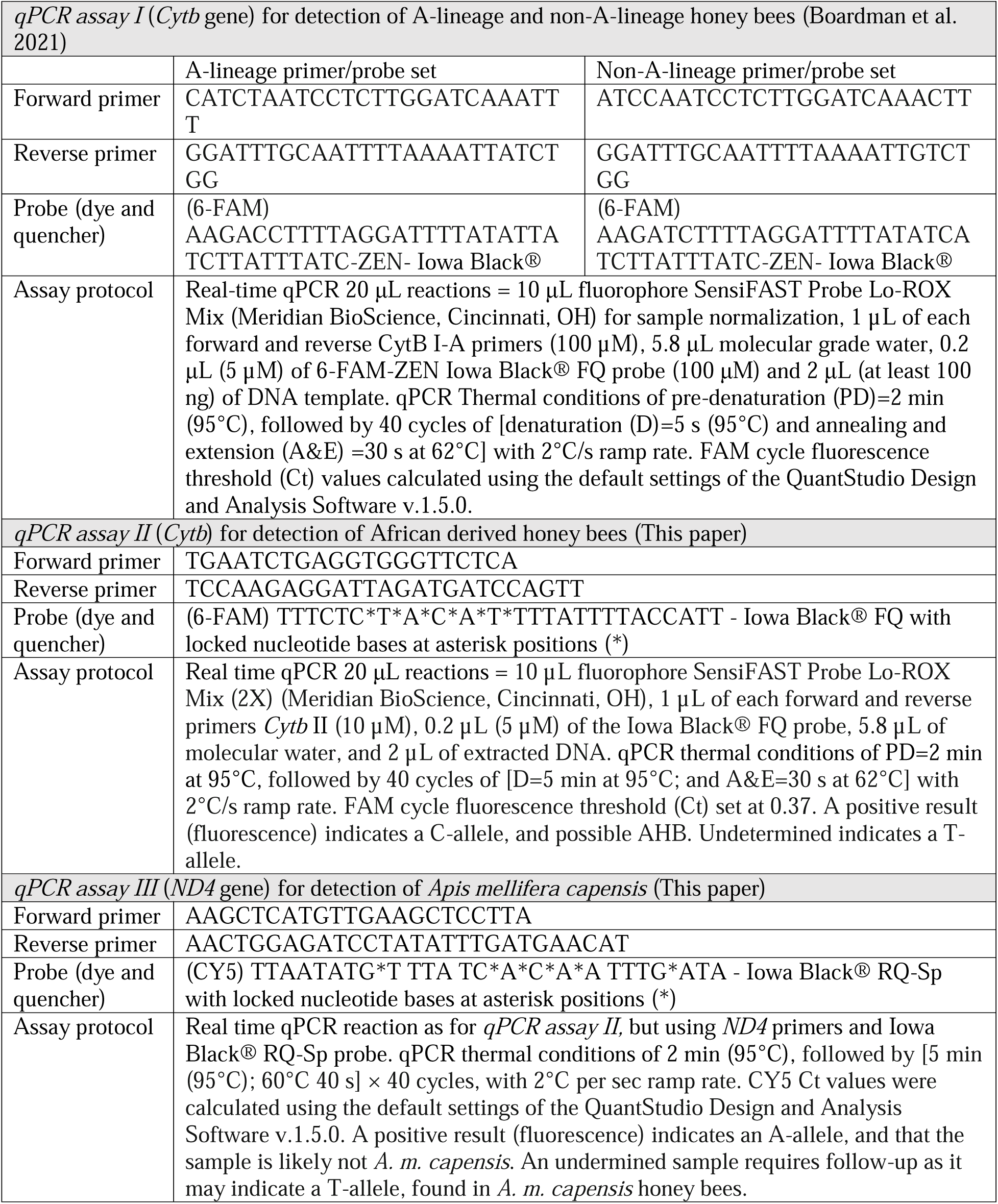
Detailed information for primers and probes and assay protocols for each of the different real-time qPCR assays. All sequences are provided in the 5’ to 3’ direction. Sensitivity analysis for the new qPCR assays are provided in Figure S5 and S6, and Table S2 and S3.

#### 2.2.2. New qPCR assay III – A. m. capensis

*qPCR assay III* amplifies a region of *ND4* and fluorescently detects an internal SNP discriminating *A. m. capensis* from all other honey bees (Figure 1) using a dual-labeled LNA probe. To develop this assay, a *NADH-dehydrogenase subunit 4* (*ND4*) sequence gene alignment was generated by extracting this gene region from the dataset of 91 mitochondrial genomes (Boardman et al., in preparation). This alignment showed four SNPs associated with a recovered *A. m. capensis* clade. One of these SNPs fell within the region previously PCR amplified, and Sanger sequenced in *Apis* species (Leelamanit et al. 2004). The alignment covering this region was extracted, and primers and probes were designed as in section 2.2.1. with small modifications. We used LNAs because there were repetitive nucleotide bases (low complexity) and to raise the annealing temperatures for probes sufficiently above the primers. Full information on primers and probes can be found in Table 2. A Ct value (fluorescence) indicates an A-allele, and that the sample is likely not *A. m. capensis*. An undetermined Ct requires follow-up as it may indicate a T-allele, found in *A. m. capensis*.

#### 2.2.3. New RFLP assay – ACMS clade

This assay was designed using 322 sequences from the NCBI GenBank nucleotide database and unpublished mitochondrial genomes (Boardman et al., in preparation). After alignment, gene sequences were converted to a BED format (*.bed) using Plink v1.9 (Purcell et al. 2007) and a phenotype file was generated, segregating *A. m. adansonii, A. m. capensis, A. m. monticola, A. m. scutellata* (ACMS clade) as cases, and all other *A. mellifera* ssp. as controls.

Following dataset pruning in Plink v1.9 for rare variants (--maf 0.05) and missing genotypes (--geno 0.05), a case-control association analysis was performed using the --assoc and –adjust qq-plot parameters for further analysis. The case/control association resulted in a significant variant located within *ND2* (raw *p* = 8.508 × 10^-59^). Given the significant segregation between the ACMS clade and all other *A. mellifera* ssp. genotypes, we further pursued genotyping of this variant in the entire dataset.

Primers for targeted genotyping were designed based on a consensus sequence obtained through NCBI Multiple Sequence Alignment Viewer using all *A. mellifera* ssp. *ND2* sequences.

Primer3 (Rozen and Skaletsky 2000) was used to design primers targeting *ND2*, with a total genomic product input of 1004 base pairs. The forward and reverse primers (Table 3) resulted in an *in silico* product spanning 646 bases. We confirmed the specificity of the selected primers using BLAST, resulting in 100% primer specificity to *A. mellifera* for both forward and reverse primers. Oligonucleotide primers were synthesized by Eurofins Genomics Inc. (Table 3). A restriction length fragment polymorphism (RLFP) genotyping assay was designed with the *in silico* primer product and target variant as input on DesignSignatures (Wright and Vetsigian 2016) using standard default tool parameters. Recommended enzymes were further evaluated with NEBCutter (Posfai et al. 2022) for target specificity, as well as optimal enzyme parameters. The enzyme PacI enzyme (5’…TTAAT^TAA…3’) (New England BioLabs Inc., Ipswich, MA, USA) was chosen for the RFLP assay based on best performance discriminating the ACMS clade from all other *A. mellifera* ssp., by discriminating between a C-allele at position 282 in the PCR gene region (ACMS), vs T-allele (other subspecies).

**Table 3.**
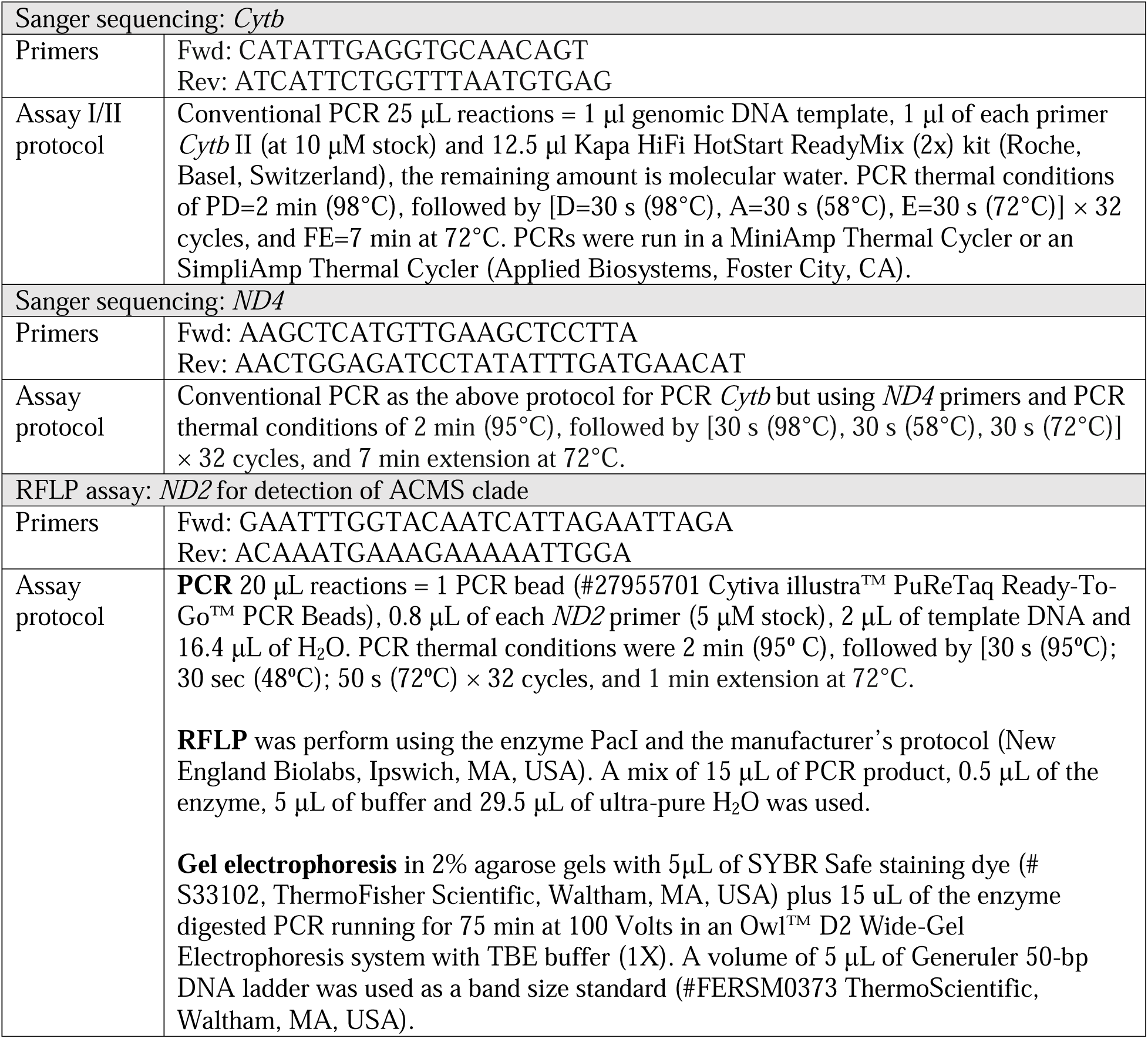
Primer information and assay protocols for conventional PCRs for *cytochrome b* (*Cytb*) and *NADH dehydrogenase 4* (ND4) genes used for Sanger sequencing to confirm qPCR results, and for RFLP assay on *NADH dehydrogenase 2* (ND2) for detection of the ACMS clade (*Apis mellifera adansonii*, *capensis*, *monticola*, and *scutellata*).

*Apis mellifera capensis* and *A. m. scutellata* (two of the regulated subspecies) have the C-allele in the targeted region of *ND2* gene (5’…TTAATCAA…3’). PacI does not cut in the region; therefore, we would expect to see only the uncut PCR product on the 2% agarose gel (Figure S3). Note that some ACMS have a PacI recognition site at 28-35bp. This may result in a cut that yields products of approximately 32bp and 614bp. However, this 32bp fragment was too small to detect on our gels and not included in our interpretation (Figure S3). In contrast, non-ACMS *A. mellifera* subspecies are predicted to have a T-allele at position 282. This creates a recognition site for PacI (5’…TTAAT^TAA…3’) resulting in a cut in the PCR product (646bp) into 2 bands of 281bp and 365bp for *A. m. intermissa* Maa, *A. m. lamarckii* Cockerell, and *A. m. jemenitica* Ruttner. Subspecies *A. m. ligustica*, *A. m. mellifera* and *A. m. meda* Skorikov also have a second PacI site at 28-35bp which predicts 3 bands: 32bp, 251bp, and 365bp. While the 32bp fragment was too small to be kept on our gel visualization, the incomplete cutting of the 281bp band into 32bp and 251bp can result in three bands of 251bp, 281bp and 365bp being visible for these subspecies (Figure S4).

In summary, subspecies from the ACMS clade (*A. m. adansonii, A. m. capensis, A. m. monticola, A. m. scutellata*) only have PCR product visible on the gel, while non-ACMS *A. mellifera* subspecies will have 2-3 bands visible after PacI digest (Figure S4).

### 2.3. Samples

To develop this workflow, we assembled a comprehensive sample set of 36 genomic DNA samples from a previous dataset (Boardman et al. 2021), and 109 adult worker honey bees and drone pupae preserved in 95% ethanol. Collectively, these samples represented 16 *A. mellifera* subspecies, and defensive and gentle honey bees from USA. Summary information for all samples can be found in Table 1, with full details provided in Table S1.

The genomic samples were 15 historical reference specimens loaned from the Ruttner Bee Collection at the Bee Research Institute, Oberursel, Germany, and corresponding to fourteen *A*. *mellifera* subspecies across the known lineages of *A. mellifera*; 18 samples from South Africa (*A. m. capensis* and *A. m. scutellata*); and three samples of immature queens from Florida, USA. These genomic samples were characterized using *qPCR assay I* and had their Cytb sequenced in a previous study (Boardman et al. 2021). Here, we ran those same samples through the newly developed assays and sequenced ND4.

The 109 new honey bee samples consisted of an additional 35 samples from South Africa (*A. m. capensis*, *A. m. scutellata* and hybrid specimens); 37 FDACS-DPI samples from hives in Florida with a defensive phenotype reported in stinging incidents (n=37, assayed blind); samples from highly defensive swarms from the USA (with 3-4 bees per colony, total samples: Arizona: n=16; Texas: n=6); bees from the USA with a gentle phenotype (Florida: n=3; Puerto Rico: n=6 - thought to display the gentle African phenotype and European ancestry *via* hybridization (Avalos et al. 2017)); one feral, defensive sample from north Florida; four bees from Florida previously identified by Fast Africanized Bee Identification System (FABIS) and/or USDA-ID (Rinderer et al. 1986, 1993; Sylvester and Rinderer 1987; Rinderer 1998); and one positive control consisting of a pooled bee sample of three honey bee workers from a colony housed at in the UF Honey Bee Research and Extension Laboratory (HBREL) with European ancestry (C-lineage).

Thus, the final dataset consisted of 53 samples from hives in 19 geographic areas of South Africa, 29 of these were assayed blind, and included one to three specimens per area. All samples are from different apiaries and colonies, except those from Pretoria and Upington.

Samples from the same colonies have previously been characterized with morphometrics, microsatellites, SNPs and mitochondrial genomes as *A. m. scutellata*, *A. m. capensis* or hybrids of the two subspecies (Eimanifar et al. 2018a, 2018b, 2020; Bustamante et al. 2020). Bee samples that were from South African colonies consistently assigned to *A. m. scutellata* and *A. m. capensis* in these studies served as positive controls for these subspecies.

### 2.4. DNA extraction

Adult honey bee worker specimens were cut longitudinally from head to thorax with a single use sterile scalpel and petri-dish to expose soft tissue. The abdomen was discarded. The head and thorax were subsequently homogenized at 4 M/s for 10s using a Benchtop FastPrep-24™ Homogenizer (MP Biomedicals LLC, Santa Ana, CA, USA, or a Mini-Beadbeater-16 (Biospec Products, Bartlesville, OK, USA). Genomic DNA extraction was completed using the DNeasy® Blood and tissue kit (#69504, Qiagen, Valencia, CA, USA) recommended protocol for Purification of Total DNA from Animal Tissues (Spin-Column Protocol). DNA yield was assessed with a Qubit (Qubit 3.0 Fluorometer, ThermoFisher Scientific, Waltham, MA) or a NanoDrop 2000 spectrophotometer (ThermoFisher Scientific, Waltham, MA). For pupae, we followed the manufacturer’s protocol without the preliminary excision step. Final genomic DNA elution was in UltraPure H_2_O.

### 2.5. qPCR assays

The *qPCR assay I* (*Cytb*) was run on 132 samples, the *qPCR assay II* (*Cytb* gene) was run on 145 samples, and the *qPCR III* (*ND4*) was run on 145 samples (Table 1). Real-time *qPCR assays I, II and III* were performed in an Applied Biosystems QuantStudio 5 platform (Table 2). Data were validated by comparing the results of qPCR with Sanger sequences (Table 3).

### 2.6. Sanger sequencing: Cytb and ND4 genes

To validate qPCR assays, and detect false positives, conventional PCRs were performed as listed in Table 3. PCR amplicon products were analyzed by 1.5% agarose gel electrophoresis. PCR products of the target size (i.e., 370bp for *Cytb* and 409bp for *ND4*) were purified using a Qiagen QIAquick PCR Purification Kit (Qiagen, Hilden, Germany). Purified PCR products were sequenced bidirectionally using a BigDye Terminator (v.3.1) cycle sequencing kit (Applied Biosystems, Foster City, CA). Sequencing was performed on an Applied Biosystems SeqStudio platform at FDACS-DPI. Contig chromatograms obtained by sequencing were analyzed using Sequencher 5.4.6 (Gene Codes Corp., Ann Arbor, MI) and assembled into sequence contigs (Gene Codes Corp., Ann Arbor, MI).

### 2.7. RFLP assay (*ND2* gene)

The *ND2* gene was amplified using conventional PCR, before digestion of PCR product with PacI (Table 3). This assay was performed on genomic DNA from honey bee samples in regions of the USA where AHBs are thought to be well established, as well as regions in South Africa to serve as positive controls for the assay (Table 1 & S1). Mitotypes were assigned as (1) ACMS clade if no cuts were observed, and (2) all other honey bees if two-three bands were observed (enzyme cuts). We observed that even using overnight incubation, incomplete digestion of the initial PCR product was observed in all experiments, and uncut PCR product of the size of the target gene fragment was present, matching expected PCR product sizes.

## 3. RESULTS

### 3.1. qPCR assay I (*Cytb* gene) - Detect A-lineage and non-A-lineage

Of the 132 honey bees tested in this bioassay, 102 samples were found to be mitochondrially A-lineage (Table 4 & S1). These A-lineage bees included 47 samples from Africa (43 from South Africa, and one sample each from Ethiopia, Kenya, Central African Republic, Morocco); 54 from the USA (Arizona, Florida, and Texas); and one sample from Europe (Malta). Thirty honey bees were found to be non-A-lineage. These bees included samples from north Africa (Egypt, Morocco), Western Asia (Syria, Turkey, Turkmenistan), and Europe (Croatia, Italy, Norway, Spain); and 21 samples from the USA (Arizona, Florida, Puerto Rico and Texas). All samples that were positive for the A-lineage assay had “=undetermined” FAM Ct values for the non-A-lineage assay, and *vice versa*.

**Table 4.**
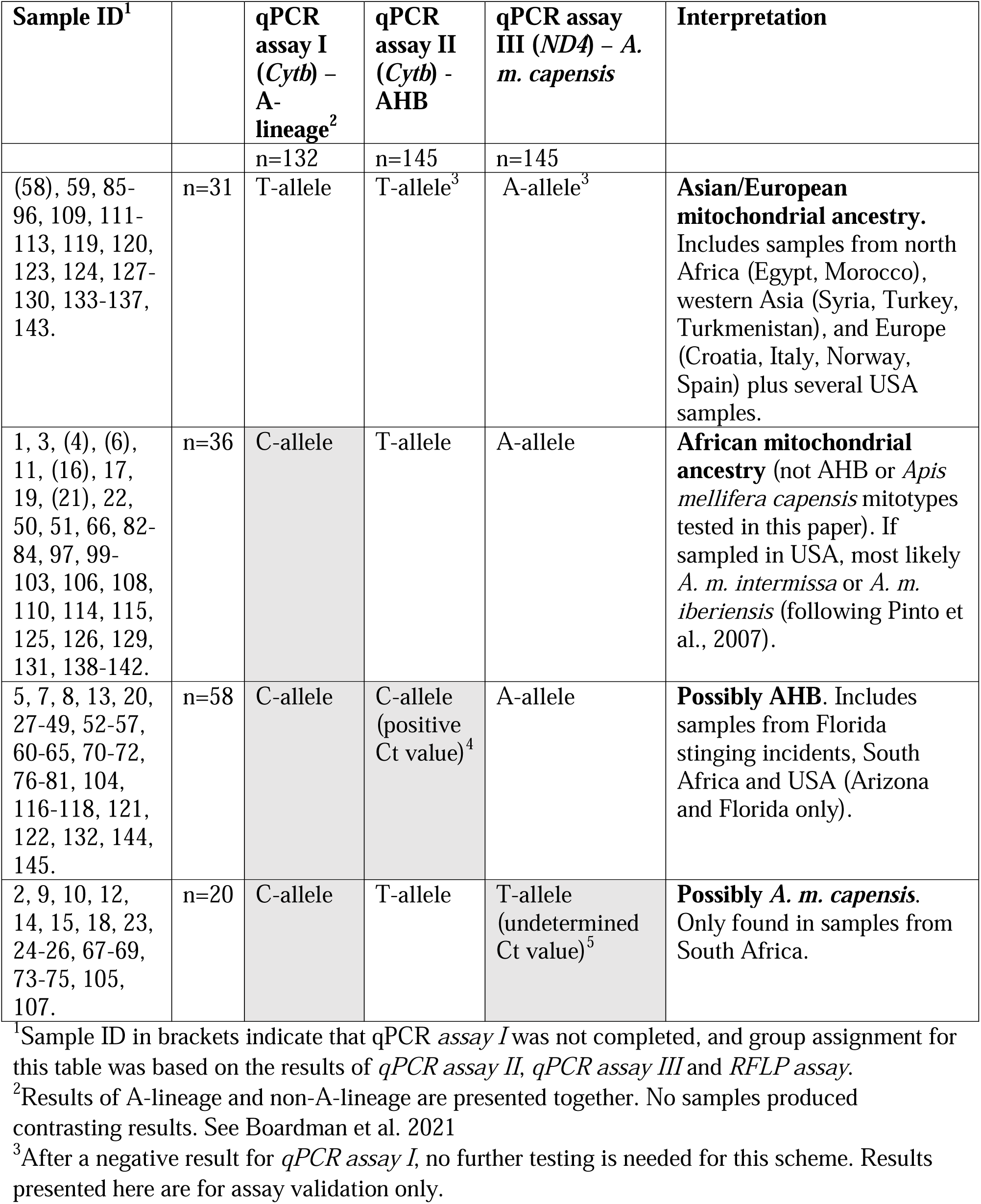
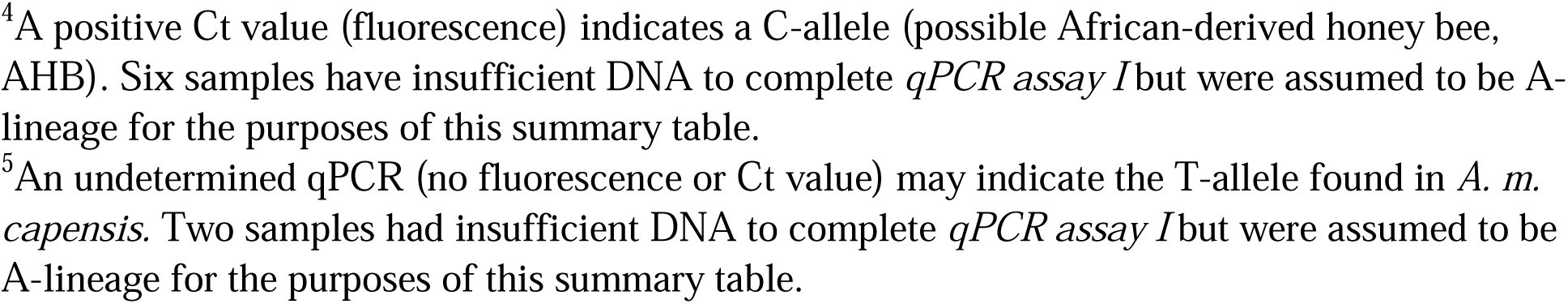
Summary of interpretation of qPCR assay results for unknown *Apis mellifera* samples.

| Sample ID<sup>1</sup> |  | qPCR assay I (*Cytb*) – A-lineage<sup>2</sup> | qPCR assay II (*Cytb*) – AHB | qPCR assay III (*ND4*) – *A. m. capensis* | Interpretation |
| --- | --- | --- | --- | --- | --- |
|  |  | n=132 | n=145 | n=145 |  |
| (58), 59, 85-96, 109, 111-113, 119, 120, 123, 124, 127-130, 133-137, 143. | n=31 | T-allele | T-allele<sup>3</sup> | A-allele<sup>3</sup> | **Asian/European mitochondrial ancestry.** Includes samples from north Africa (Egypt, Morocco), western Asia (Syria, Turkey, Turkmenistan), and Europe (Croatia, Italy, Norway, Spain) plus several USA samples. |
| 1, 3, (4), (6), 11, (16), 17, 19, (21), 22, 50, 51, 66, 82-84, 97, 99-103, 106, 108, 110, 114, 115, 125, 126, 129, 131, 138-142. | n=36 | C-allele | T-allele | A-allele | **African mitochondrial ancestry** (not AHB or *Apis mellifera capensis* mitotypes tested in this paper). If sampled in USA, most likely *A. m. intermissa* or *A. m. iberiensis* (following Pinto et al., 2007). |
| 5, 7, 8, 13, 20, 27-49, 52-57, 60-65, 70-72, 76-81, 104, 116-118, 121, 122, 132, 144, 145. | n=58 | C-allele | C-allele (positive Ct value)<sup>4</sup> | A-allele | **Possibly AHB.** Includes samples from Florida stinging incidents, South Africa and USA (Arizona and Florida only). |
| 2, 9, 10, 12, 14, 15, 18, 23, 24-26, 67-69, 73-75, 105, 107. | n=20 | C-allele | T-allele | T-allele (undetermined Ct value)<sup>5</sup> | **Possibly *A. m. capensis*.** Only found in samples from South Africa. |

<sup>1</sup>Sample ID in brackets indicate that qPCR assay I was not completed, and group assignment for this table was based on the results of qPCR assay II, qPCR assay III and RFLP assay.
<sup>2</sup>Results of A-lineage and non-A-lineage are presented together. No samples produced contrasting results. See Boardman et al. 2021
<sup>3</sup>After a negative result for qPCR assay I, no further testing is needed for this scheme. Results presented here are for assay validation only.
<sup>4</sup>A positive Ct value (fluorescence) indicates a C-allele (possible African-derived honey bee, AHB). Six samples have insufficient DNA to complete *qPCR assay I* but were assumed to be A-lineage for the purposes of this summary table.
<sup>5</sup>An undetermined qPCR (no fluorescence or Ct value) may indicate the T-allele found in *A. m. capensis*. Two samples had insufficient DNA to complete *qPCR assay I* but were assumed to be A-lineage for the purposes of this summary table.

### 3.2. qPCR assay II (*Cytb* gene) – Detect AHB

The positive samples for this *qPCR assay II* included 32 FDACS-DPI honey bee specimens from Florida, USA that were collected from stinging incidents or public complaints, providing a correlation between genetic data and behavioral data. An additional 26 samples were positive for the *qPCR assay II* marker for this SNP. These included 18 samples from South Africa and 8 samples from the USA (Arizona, n=3; and Florida, n=5) (Table 4). This SNP was not detected in samples from Puerto Rico or Texas, USA. Sanger sequencing confirmed the results of the qPCR for all samples, with positive FAM Ct values indicating a “C”, and undetermined qPCR assays indicating a T-allele. All new *CytB* sequencing data have been deposited in GenBank (accession numbers: OQ675833 – OQ675976).

### 3.3. qPCR assay III (ND4 gene) – Detect possible A. m. capensis samples

Twenty samples had undetermined FAM Ct values for this marker (T-allele), suggesting a possible *A. m. capensis* identity (Table 4). All these samples were from South Africa and were exclusively from locations in regions of South Africa known to have pure or hybrid populations of *A. m. capensis* (Hepburn et al. 1994), conceivably with *A. m. capensis* maternal genes. The remaining 125 other *A. mellifera* samples returned positive FAM Ct values indicating that they are likely other *A. mellifera* subspecies. Sanger sequencing confirmed that qPCR negative samples had a T-allele (possible *A. m. capensis*), while samples with positive Ct values had an A-allele. The sequencing revealed one false positive for the qPCR assay. The sample was *A. m. litorea* Smith from South Africa (a subspecies found in narrow region along the east coast of Africa). The qPCR returned a negative qPCR result, while the Sanger sequencing showed an A-allele – contrasting with the qPCR result. The qPCR was repeated to check the result, which was confirmed. All *ND4* sequences have been deposited into GenBank (accession numbers: OQ675977 – OQ676119).

### 3.4. RFLP assay (*ND2* gene) – Detect ACMS clade

We were able to obtain results for 142 samples. One A-lineage bee and two non-A-lineage bees failed to amplify for this RFLP (Table S1). Of the 142 results, 105 samples could be assigned to the ACMS clade (Table S1) indicating they have a C-allele, which results a mismatch in the restriction enzyme recognition site, and thus the PCR product remains uncut (i.e., only the uncut PCR product (ca. 646bp) was visible on the gel (Figure 2). Ninety-three bees previously identified as A-lineage with our *qPCR assay I* (*Cytb*) marker could be assigned to the ACMS clade by RFLP testing, and the remaining 12 ACMS samples were samples that were not tested using *qPCR assay I*. One of these samples was from Florida, while the remaining 11 were from South Africa.

**Figure 2.**
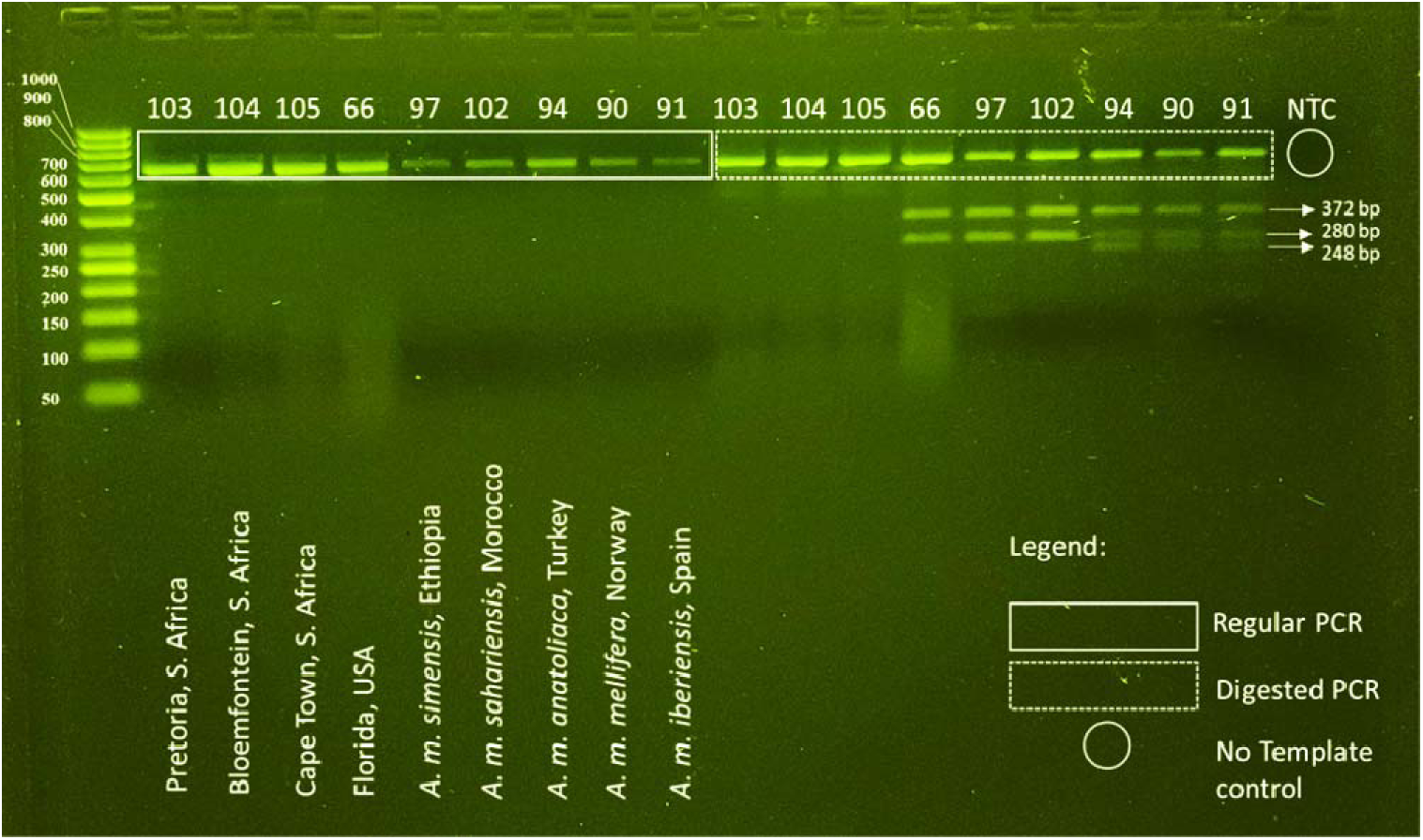
Representative RFLP gel band patterns of PacI digest on *NADH dehydrogenase 2* (*ND2*) gene with *Apis mellifera* honey bee specimens of African descent *vs.* other origins. Lane numbers refer to honey bee specimen numbers as listed in Table 1 & S1. Lanes 1 to 9 correspond to 5 μL regular PCR products, while lanes 10 to 18 show 15 μL digested PCR with enzyme PacI. No template control (NTC) is shown in lane 19. Undigested PCR products are visible in all lanes and were excluded for discrimination purposes. Figure S3 shows the *in silico* predictions, and Figure S4 shows the same PCR product, but 35 μL digested PCR.

The 37 samples that were not ACMS clade (the restriction enzyme cutting leading to visible bands on the gel, indicative of a T-allele), consisted of 28 non-A-lineage, 8 A-lineage, and one sample that was not tested in *qPCR assay I*. While the exact number of bands is not relevant for this assay to be effective as ACMS samples will not be cut by PacI, we did notice differences in the banding pattern of non-ACMS samples. The three-band pattern that we would expect to find in non-A-lineage samples commonly found in Europe and western Asian subspecies (e.g., *A. m. ligustica*, *A. m. mellifera* and *A. m. meda*) was found in 26 of the 28 non-A-lineage samples (e.g., Figure 2). The non-A-lineage historical reference samples of *A. m. lamarckii* (Sample ID: 89) and *A. m. syriaca* Skorikov (Sample ID: 88) had two-bands.

Of the eight A-lineage samples that were found not to be in the ACMS clade, four samples had the three-band pattern, and four samples had the two-band pattern. The samples with the three-band pattern were the historical reference sample *A. m. ruttneri* (Sample ID: 99), and three samples from one colony in Texas, USA (Sample ID: 110, 125, 126). The two-band pattern samples were *A. m. sahariensis* Baldensperger (Sample ID: 97), *A. m. simensis* Meixner, Leta, Koeniger and Fuchs (Sample ID: 102), one sample from South Africa (Sample ID: 18) and one from Florida, USA (Sample ID: 66).

No non-A-lineage sample from *qPCR assay I* was assigned to the ACMS clade, further reinforcing the assessment of their maternal ancestry as from non-A-lineage subspecies.

## 4. DISCUSSION

The United States Animal and Plant Inspection Services (APHIS) data from the period of 2001-2021 shows that honey bees (*Apis* sp.) are regularly intercepted at air, maritime and terrestrial ports within the USA. Such intercepts account for 82.9% of all Apoidea intercepts (Marcelino et al. 2022). There is a need to detect these organisms expediently and accurately at ports of entry within the continental USA and/or prior to importation, and for apiary inspection and queen breeding certification. Here, we developed a series of qPCR assays to establish the maternal ancestry of an unknown honey bee using clade-specific and/or subspecies-specific assays based on a single SNP. These qPCR assays were able to identify one mitotype of AHB (correlated with defensive behaviors in Florida), and at least one mitotype of *A. m. capensis*. We also developed an RFLP assay for differentiating *A. m. adansonii, A. m. capensis, A. m. monticola*, *A. m. scutellata* ancestry (ACMS clade) from all other *A. mellifera* samples. These assays represent useful progress in developing more informative markers, while being cognizant of developing methods that can be widely used by researchers with varying levels of resources. As part of our marker validation, we also generated additional single gene sequences of mitochondrial *Cytb* and *ND4* and uploaded those to GenBank. They can now be used as additional references for future work.

The existing A-lineage and non-A-lineage assays (Boardman et al. 2021) are useful as an initial qPCR, however, they cannot further discern the A-lineage. Our data suggests that it would be satisfactory to run either one of these qPCRs as the results are congruent. No samples classified as non-A-lineage were found to be positive for our AHB or *A. m. capensis* markers.

Any honey bees assigned as non-A-lineage using this assay should have Asian or European maternal ancestry.

Our new *Cytb* marker (*qPCR assay II*) was able to detect defensive AHBs in Florida and some from Arizona, while other suspected AHBs were negative, and likely have other maternal ancestry. The bees that were taken to Brazil and subsequently disseminated across the Americas are from Klipfontein, a region of South Africa north of Pretoria (Kerr 1992) with high mitotype diversity (Moritz et al. 1994). Indeed, previous studies have shown variation in mitotypes for AHBs present in the USA (e.g., Szalanski and McKern 2007). Furthermore, not all honey bees suspected of being AHBs possess the mitochondrial DNA SNPs consistent with *A. m. scutellata*-derived from southern Africa regions. For example, suspected AHBs from southern Texas and New Mexico matched mitotypes common in north African *A. m. intermissa* or Portuguese and Spanish *A. m. iberiensis* Engel (Pinto et al. 2007). The SNP marker targeted in this paper can be found in some samples from the known invasive range of Mexico, (KJ601784, Gibson and Hunt 2016), Brazil (EF184030.1, Pinto et al. 2007), USA (EF016648, Szalanski and McKern 2007), and Mbo-A/Hinf-1(1) haplotype (EU513284, Ferreira et al. 2009). This suggests that the marker traces one mitotype that originated in South Africa and moved through Brazil, into the USA.

We included four samples that had been previously identified as African/Africanized using FABIS, and three of which were also identified using USDA-ID (Table S1). Our data confirmed the findings of FABIS/USDA-ID in two samples. The positive control consisting of honey bees from the HBREL apiary and assumed to be of European ancestry returned as non-A-lineage, suggesting European mitochondrial ancestry. This mixture of consensus and contrasting results is not surprising, given that FABIS and USDA-ID classify honey bees morphometrically, a process that has inherent variability and that can fail to find low levels of AHB influence in a colony (Guzmán-Novoa and Page 1994; Guzmán-Novoa et al. 1994), while we used molecular markers to identify the honey bees.

Though our molecular marker predicted African ancestry, we are not suggesting that this marker causes defensive behavior. Defensive behavior in *A. mellifera* has been attributed to several different alleles and is not determined only by African matrilines (Avalos et al. 2017; Harpur et al. 2020). In our data, 32 of the 37 honey bee samples collected from Florida reported as stinging incidents were positive for our AHB marker. In addition, three samples from one colony from Arizona were positive for the AHB marker. In contrast, other samples from Arizona, and Texas were negative, despite having a defensive phenotype (see additional detailed discussion in supplementary materials). One sample from north Florida (feral, defensive) was also negative (Sample ID: 119). Defensive behavior can be displayed by honey bees with both African and non-African maternal ancestry and is not a reliable phenotype to assign ancestry.

While our marker is not positive for all suspected AHB, it can serve as an additional important diagnostic tool for apiary inspectors/diagnosticians in Florida.

A gentle phenotype of AHB, denoted gAHB, can be found in Puerto Rico and has been previously found to be derived from AHBs in Texas (Galindo-Cardona et al. 2013; Acevedo Gonzalez et al. 2019). We tested three samples of gAHB and three from a defensive colony in Puerto Rico, and none were positive for our AHB marker. All six samples were determined to be non-A-lineage, suggesting maternal ancestry from an Asian or European subspecies. Using whole genome SNPs, Acevedo-Gonzalez et al. (2019) found that samples of gAHB in Puerto Rico formed a clade with samples from Argentina and AHBs from Texas, while samples from Arizona, AHB Brazil and A-group Africa formed a sister clade (Acevedo Gonzalez et al. 2019). Samples that were positive for A-lineage, but not positive for AHB, may have maternal ancestry from one of several different A-lineage subspecies (especially north African subspecies known to be present in USA, see discussions in Pinto et al. 2007; Carpenter and Harpur 2021), or alternatively may be *A.m. scutellata* derived from a different matriline.

The original *A. m. scutellata* introduced into Brazil were queens (Kerr 1957), thus all their offspring should, theoretically, have the same mitotypes (see discussions in Schneider et al. 2004). Given the high mitotype diversity in some regions of South Africa (Moritz et al. 1994), it is conceivable that all the queens taken to Brazil did not have the same mitotype, thus, finding a single SNP marker for all AHBs present in the USA may not be possible. Additional research into the *Cytb* mitotype targeted here would provide better understanding of the dissemination of AHBs through the Americas.

The new *qPCR assay III* targeting a *ND4* marker shows potential for *A. m. capensis* identification, possibly of the mitotype associated with *A. m. capensis* in the southwestern region of South Africa (for discussions on this, see Moritz et al. 1994; Hepburn et al. 1998; Hepburn and Radloff 2002). Of the 34 presumed *A. m. capensis* samples (based on collection locations), 19 of these (∼56%) were positive for our marker. Perhaps most significant, none of the 19 presumed *A. m. scutellata* samples have the marker. Identification of bees from South Africa as either *A. m. capensis* or *A. m. scutellata* can be difficult due to the occurrence of a wide hybridization zone. Recent papers looking at these subspecies in South Africa have shown that even phenotypes utilized in morphometric discrimination consist of a spectrum that does not follow clear geographical limitations (Eimanifar et al. 2018a, 2018b; Bustamante et al. 2020; Patterson Rosa et al. 2021). Here, our developed *qPCR assay III* serves as a method to identify the SNP in *ND4* found in a clade containing only *A. m. capensis* from regions of South Africa known to be endemic to *A. m. capensis*. Samples from three locations were always *A. m. capensis* in our assays. These locations were Cape Town, Riversdale and Touwsrivier; while several nearby locations had 1-2 samples positive for *A. m. capensis* marker, and others negative. The one sample that gave us a false positive for this qPCR assay III for *A. m. capensis* was *A. m. litorea*, a subspecies found along the east coast of Africa, stretching from Kenya to the Mozambique-South Africa border (Ruttner 1988). There are known similarities between *A. m. litorea* and *A. m. capensis* (color, morphometrics, ovariole numbers, even spermatheca), and further genetic work on this subspecies would be revealing.

We have no way to discern from our data if the honey bees positive for this SNP would have exhibited the thelytoky phenotype common among *A. m. capensis* (Onions 1914).

Nevertheless, the *A. m. capensis* marker developed here is indicative of the subspecies or introgression with *A. m. scutellata*, both of which are bees of regulatory concern in the USA. We hope this marker serves as a starting point for further research into *A. m. capensis* mitotypes, and its utility as a potential marker for this honey bee.

If we consider the 53 samples studied from South Africa (excluding historical samples from Oberursel), 37 could be assigned to AHB (*A.m. scutellata* in this case, n=18) or *A.m. capensis* (n=19) using our qPCR assay markers. As expected, no sample was positive for both markers. A full discussion of these samples is provided in supplementary materials. Twelve of the remaining 16 samples were A-lineage, but not AHB or *A. m. capensis*. The remaining four samples were also not AHB or *A. m. capensis*, but *qPCR assay I* was not run on these samples and so we cannot confirm their lineages (Table S1, assumed *A-lineage* based on RFLP *ND2*-PacI assay). These A-lineage honey bees may be alternative mitotypes of *A. m. capensis* or *A. m. scutellata* not detected by our assays, or alternatively, may represent *A. m. adansonii* or *A. m. litorea* samples (Moritz et al. 1994; Hepburn and Radloff 2002). These subspecies can be found in the northwest (*A. m. adansonii*) and northeast (*A. m. litorea*) regions of South Africa. Less likely, but still plausible, queens from A-lineage honey bees may have been imported from other regions of Africa (Hepburn 1990).

Our RFLP *ND2*-PacI assay could serve as an alternative for detecting *A. m. scutellata* and *A. m. scutellata* derived honey bees for some researchers. RFLP assays can be technically difficult to troubleshoot, with challenges regarding poor standardization and reproducibility of PCR conditions. However, the RFLP *ND2* assay we present here could serve as a viable alternative for individuals wishing to identify *A. m. scutellata* or *A. m. capensis* honey bees but who lack access to qPCR technology. While it cannot discern between each of these subspecies or *A. m. adansonii* or *A. m. monticola*, it can still be useful. *Apis mellifera adansonii* has been reported as being defensive (Kasangaki et al. 2018), and *A. m. monticola* has a different phenotype, being a larger, darker colored honey bee found only in mountainous regions of Africa. As *A. m. capensis* is also a subspecies of regulatory concern due to the thelytoky phenotype, at least in a US context, none of these subspecies would be allowed to be introduced.

High-throughput, next-generation sequencing of targeted genes or nuclear genomes is commonly used to untangle complexes of species and subspecies (Donthu et al. 2024; Momeni et al. 2021). However, these techniques are not overly useful for quick identification of a specimen to species/subspecies because they can be costly and require highly specialized labor, technology, and bioinformatics infrastructure. The development of future markers needs to include several other African subspecies and not just compare *A. m. scutellata* or *A. m. capensis* to Asian and European subspecies. In addition, including several mitotypes would be beneficial if methods are targeting mitochondrial genes. Using mitochondrial genes for markers is advantageous as they track maternal ancestry, but can also be limited given the admixed composition of AHBs. While these methods can provide information about maternal genetic background, they are not necessarily useful for other aspects of sample biology or for determining phenotype. For example, genetic components that play a role in defensive behaviors are likely of nuclear origin rather than mitochondrial origin, as occurs with other phenotypes of interest such as ovariole number and coloration (Patterson Rosa et al, 2021). While there is still a need for additional research, the assays developed in this study offer good resolution and delineate some sub-populations within the A-lineage.

## Acknowledgements

We graciously thank beekeepers in the United States of America and South Africa, members of the University of Florida Honey Bee Research and Extension Laboratory, and Florida apiary inspectors (Florida Department of Agriculture and Consumer Services – Division of Plant Industry) for providing samples. We thank Dr. Vanessa Mathieu-Sheltry (New England Biolabs, Ipswich, MA, USA) for guidance using and optimization of endonucleases. We also thank Drs. Rebecca Kimball and Edward Braun (both University of Florida) for help with *Cytb* and *ND2* alignments and trees that facilitated this project.

## Statements and declarations

### Author contributions

The research idea was conceived by James Ellis and Leigh Boardman. Samples were obtained by Jose Marcelino, Matthew Moore, Bernd Grünewald, Mike Allsopp, Christian Pirk, Garth Cambray, and James Ellis. Assay design and data collection were completed by Jose Marcelino, Leigh Boardman, Matthew Moore, James Fulton, Hector Urbina, and Laura Patterson Rosa. The first draft of the manuscript was written by Jose Marcelino and Leigh Boardman and all authors commented on successive versions of the manuscript. All authors read and approved of the final manuscript.

### Ethics approval

No ethics approval was required.

### Data availability

All data are available within the paper and its Supplementary Information. Sequences have been deposited into GenBank: OQ675833 – OQ676119).

