## Supplementary Materials for "Development of molecular markers for western honey bee (*Apis mellifera* L.) subspecies of regulatory concern in the United States"

**Electronic supplementary materials for Marcelino, Boardman et al.**

This document contains the following:

1. Dataset information (Data S1, Data S2)
   a. Data S1 and S2 are separate .nex files
2. Supplementary figures and tables (Figures S1-S6, Table S2-S3)
   a. Table S1 is a separate .xlsx file
3. Supplementary detailed discussion of specific results

**Datasets**

**File: Data S1 – Marcelino_Boardman_CYTB_alignment.nex**

The *cytochrome b* (*Cytb*) and *NADH dehydrogenase subunit 2* (*ND2*) alignments used to develop *qPCR assay II* to detect African-derived honey bees (AHBs, section 2.2.1), and *RFLP assay* to detect A-lineage clade that includes *Apis mellifera adansonii,* *A. m. capensis,* *A. m. monticola*, and *A. m. scutellata* (ACMS clade*,* section 2.2.3) are provided. These data are collated from NCBI Genbank (ncbi.nlm.nih.gov), from both single gene sequences, as well as the appropriate gene region extracted from mitochondrial genome submissions. Unpublished data are from full mitochondrial genomes sequences (Boardman et al., in preparation), which will be uploaded to GenBank as part of that submission.

**File: Data S2 – Marcelino_Boardman_ND2_alignment.nex**

The NADH *dehydrogenase subunit 4* (*ND4*) dataset used for *qPCR assay III – A. m. capensis* (section 2.2.2) was this gene region extracted from full mitochondrial genomes sequences (Boardman et al., in preparation), as there are fewer ND4 single gene sequences on GenBank, and no additional single gene *A. m. capensis* samples.

**Supplementary figures and tables**

**Figure S1: Phylogenetic tree of *cytochrome b* (*Cytb*) alignment data (Data S1) used to develop the *qPCR assay II* – African-derived honey bee (AHB).** GenBank accession numbers are provided for all published data. Location and author-assigned subspecies or haplotype information are provided for some data from mitochondrial reference genomes. Data that contained the informative C-allele targeted by the qPCR assay are shown inside yellow rectangles, together with location or haplotype information. All other *Apis mellifera* samples had a T-allele at this site. The tree is midpoint rooted. Node labels indicate bootstrap values, and unlabeled lineages are 100%.

**Figure S2: Phylogenetic tree of *NADH dehydrogenase subunit 2* (*ND2*) alignment data (Data S1) used to develop the RFLP assay to detect A-lineage clade that includes the honey bees *Apis mellifera adansonii*, *A. m. capensis*, *A. m. monticola*, and *A. m. scutellata* (ACMS clade).** GenBank accession numbers are provided for all published data. Location and author-assigned subspecies or haplotype information are provided for some data from mitochondrial reference genomes. Data that contained the informative C-allele that disrupts the *PacI* recognition site are shown inside the blue rectangle, together with location or haplotype information. One additional sample (KT828518) that was not thought to be part of ACMS clade also had a C-allele, and four samples had a G-allele at this site which would disrupt the restriction enzyme. These exceptions are noted in parenthesis after the accession number. All other *A. mellifera* samples had a T-allele at this site. The tree is midpoint rooted. Node labels indicate bootstrap values, and unlabeled lineages are 100%.

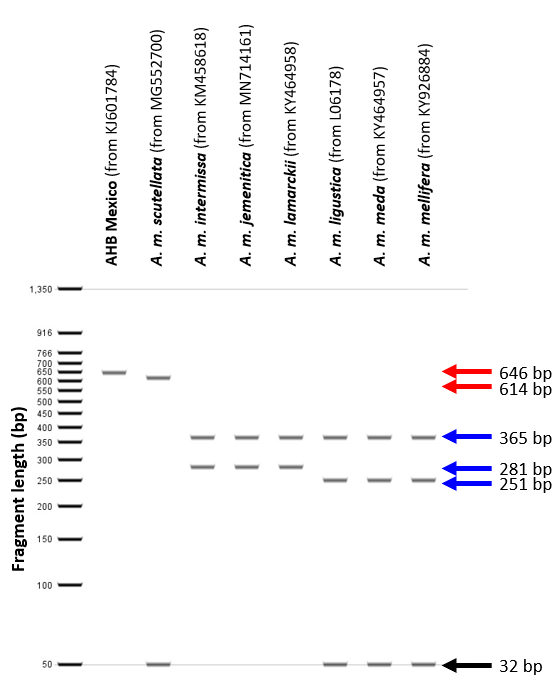

**Figure S3: *In silico* prediction for RFLP assay with PacI digest of PCR amplified region of *NADH dehydrogenase 2* (*ND2*) from mitochondrial genome sequences from Genbank.** Samples from the ACMS clade will have single 646 bp band indicating an undigested PCR product, or single 614 bp band (red arrows) if the sample has the additional recognition site that yields a 32 bp fragment. Other samples have various banding patterns (blue arrows) with the 646 bp fragment cut into 365 bp and 281 or 251 bp. The small 32 bp band (black arrow, shown here as 50 bp based on the ladder chosen) is not visible on our gels. Incomplete cutting of the 281 bp band resulted in bands of 281, 251 bp + 32 bp in our samples (see Figure 2 and S4). Generated in Geneious version 11.0.5.

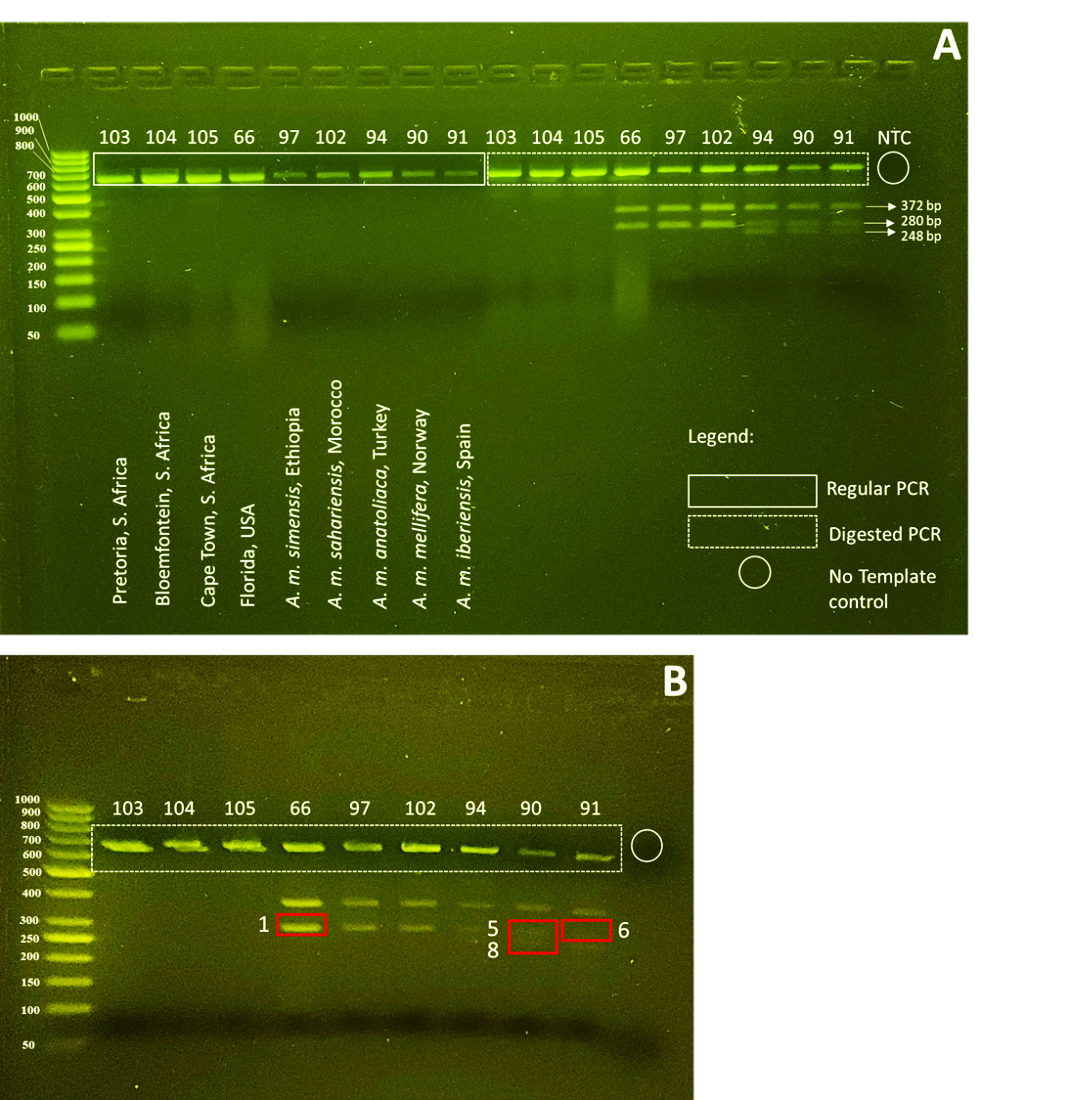

**Figure S4.** **Representative RFLP gel band patterns of PacI digest on *NADH dehydrogenase 2* (*ND2*) gene with honey bee specimens of African descent *vs.* other origins.** Lane numbers refer to honey bee specimen numbers as listed in Table 1 and Table S1, with a no template control (NTC) shown in lane 10. Samples shown here are repeated from Figure 2, but with 35 μL digested PCR with enzyme *PacI*. Bands enclosed in red squares were excised and Sanger sequenced with primers ND2_Fw 5’- GAATTTGGTACAATCATTAGAATTAGA-3’ and ND2_RV_XL 5’-TGCTTTCAATTTATTATTTCATAAG-3’ for bands 1, 5 and 6 and ND2_Fw_XS 5’-taatattaaatccacaaataa-3’ and ND2_Rv_XS 5’-attaatgttgatattaaaa-3’ for band 8.

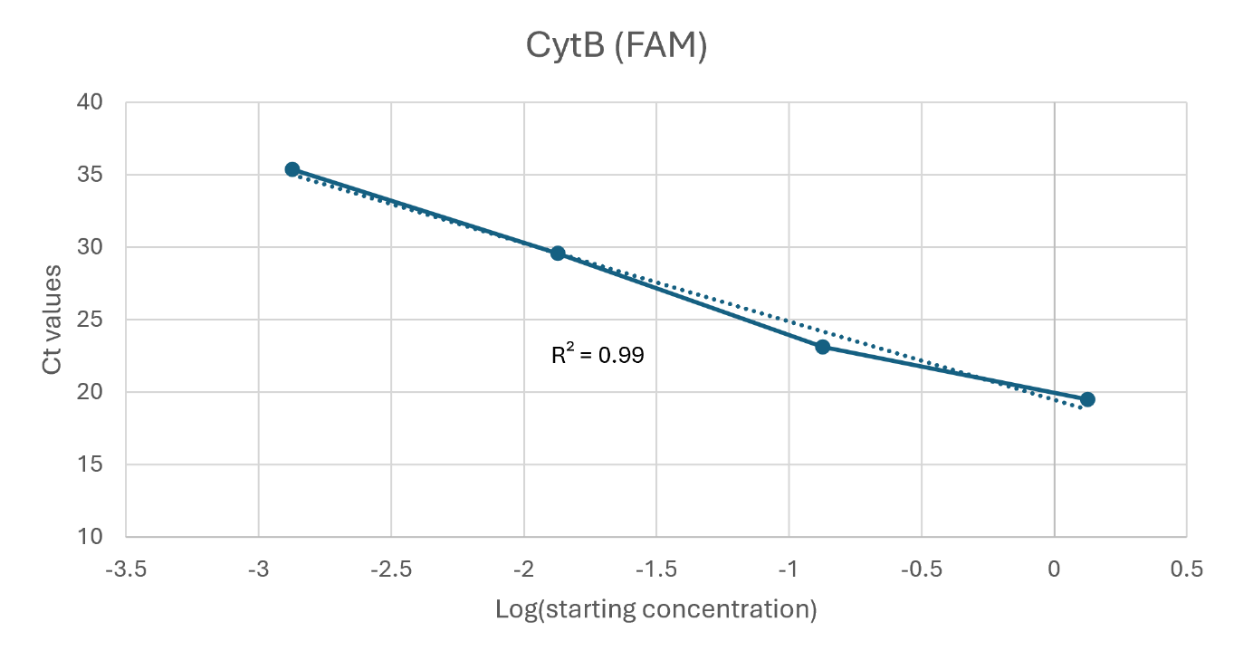

**Figure S5. PCR efficiency of qPCR assay II (*Cytb*).** Pearson coefficient shown. Positivity identified samples (based on Sanger sequencing) were used for this sensitivity analysis. The starting DNA concentration was quantified on a fluorometer (Waltham, MA) according to manufacturer’s recommended protocols. Real time qPCR reactions were run as described in Table 2 qPCR assay II (*Cytb*), and results are provided in Table S1.

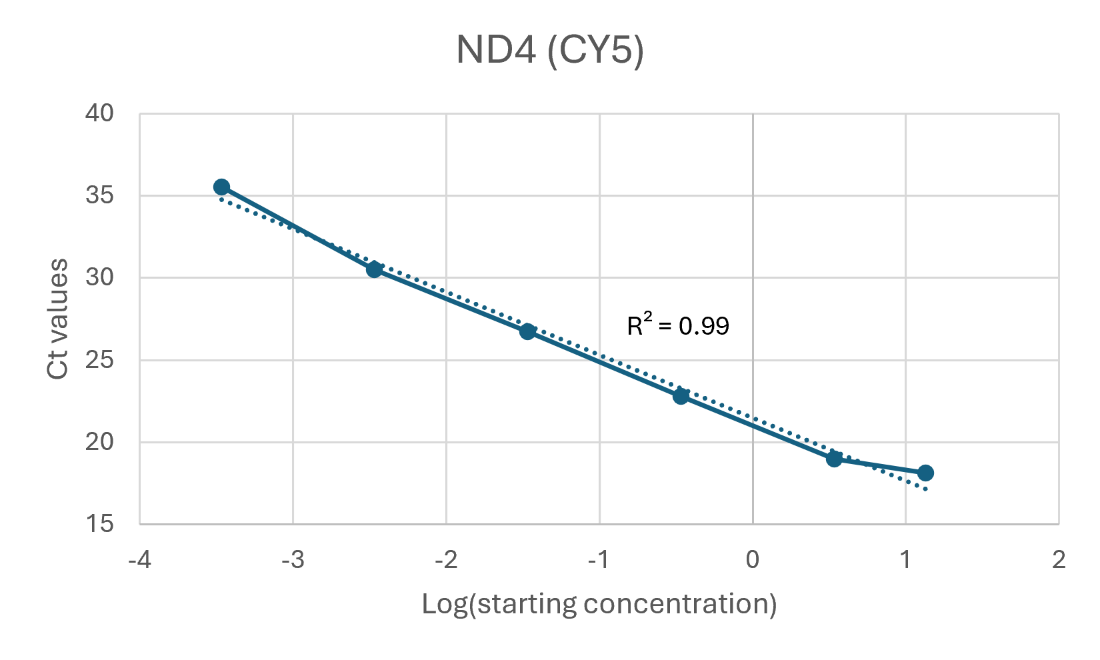

**Figure S6. PCR efficiency of qPCR assay III (ND4).** Pearson coefficient shown. Positivity identified samples (based on Sanger sequencing) were used for this sensitivity analysis. The starting DNA concentration was quantified on a fluorometer (Waltham, MA) according to manufacturer’s recommended protocols. Real time qPCR reactions were run as described in Table 2 qPCR assay III (*ND4*), and results are provided in Table S2.

**File: Table S1 - Marcelino_Boardman_Suppl.xlsx**

**Table S1. Detailed sample information and results of qPCR assays I, II, III and RFLP data discriminated by table heading color.**

**Table S2. Results of qPCR assay II (*Cytb*) sensitivity analysis.**

| **Sample Name** | **Target Name** | **Task** | **Reporter** | **Quencher** | **CT** |
| --- | --- | --- | --- | --- | --- |
| NTC | Target 1 | NTC | FAM | NFQ-MGB | Undetermined |
| NTC | Target 1 | NTC | FAM | NFQ-MGB | Undetermined |
| NTC | Target 1 | NTC | FAM | NFQ-MGB | Undetermined |
| Non-scutellata Control | Target 1 | STANDARD | FAM | NFQ-MGB | Undetermined |
| Non-scutellata Control | Target 1 | STANDARD | FAM | NFQ-MGB | Undetermined |
| Non-scutellata Control | Target 1 | STANDARD | FAM | NFQ-MGB | Undetermined |
| Scutellata Control | Target 1 | STANDARD | FAM | NFQ-MGB | 15.91 |
| Scutellata Control | Target 1 | STANDARD | FAM | NFQ-MGB | 15.975 |
| Scutellata Control | Target 1 | STANDARD | FAM | NFQ-MGB | 15.926 |
| SCU E-1 | Target 1 | STANDARD | FAM | NFQ-MGB | 19.559 |
| SCU E-1 | Target 1 | STANDARD | FAM | NFQ-MGB | 19.474 |
| SCU E-1 | Target 1 | STANDARD | FAM | NFQ-MGB | 19.481 |
| SCU E-2 | Target 1 | UNKNOWN | FAM | NFQ-MGB | 23.451 |
| SCU E-2 | Target 1 | UNKNOWN | FAM | NFQ-MGB | 22.964 |
| SCU E-2 | Target 1 | UNKNOWN | FAM | NFQ-MGB | 22.999 |
| SCU E-3 | Target 1 | UNKNOWN | FAM | NFQ-MGB | 29.395 |
| SCU E-3 | Target 1 | UNKNOWN | FAM | NFQ-MGB | 29.784 |
| SCU E-3 | Target 1 | UNKNOWN | FAM | NFQ-MGB | 29.609 |
| SCU-4 | Target 1 | UNKNOWN | FAM | NFQ-MGB | 34.145 |
| SCU-4 | Target 1 | UNKNOWN | FAM | NFQ-MGB | 35.808 |
| SCU-4 | Target 1 | UNKNOWN | FAM | NFQ-MGB | 36.148 |
| SCU E-5 | Target 1 | UNKNOWN | FAM | NFQ-MGB | Undetermined |
| SCU E-5 | Target 1 | UNKNOWN | FAM | NFQ-MGB | 37.375 |
| SCU E-5 | Target 1 | UNKNOWN | FAM | NFQ-MGB | Undetermined |
| SCU E-6 | Target 1 | UNKNOWN | FAM | NFQ-MGB | Undetermined |
| SCU E-6 | Target 1 | UNKNOWN | FAM | NFQ-MGB | Undetermined |
| SCU E-6 | Target 1 | UNKNOWN | FAM | NFQ-MGB | Undetermined |

**Table S3. Results of qPCR assay III (*ND4*) sensitivity analysis.**

| **Sample Name** | **Target Name** | **Task** | **Reporter** | **Quencher** | **CT** |
| --- | --- | --- | --- | --- | --- |
| NTC | Target 2 | NTC | CY5 | NFQ-MGB | Undetermined |
| NTC | Target 2 | NTC | CY5 | NFQ-MGB | Undetermined |
| NTC | Target 2 | NTC | CY5 | NFQ-MGB | Undetermined |
| Scutellata Control | Target 2 | STANDARD | CY5 | NFQ-MGB | 18.15062 |
| Scutellata Control | Target 2 | STANDARD | CY5 | NFQ-MGB | 18.29861 |
| Scutellata Control | Target 2 | STANDARD | CY5 | NFQ-MGB | 17.97075 |
| Non-scutellata Control | Target 2 | STANDARD | CY5 | NFQ-MGB | 19.188 |
| Non-scutellata Control | Target 2 | STANDARD | CY5 | NFQ-MGB | 18.705 |
| Non-scutellata Control | Target 2 | STANDARD | CY5 | NFQ-MGB | 19.048 |
| Non-scutellata E-1 | Target 2 | UNKNOWN | CY5 | NFQ-MGB | 22.719 |
| Non-scutellata E-1 | Target 2 | UNKNOWN | CY5 | NFQ-MGB | 23.021 |
| Non-scutellata E-1 | Target 2 | UNKNOWN | CY5 | NFQ-MGB | 22.613 |
| Non-scutellata E-2 | Target 2 | UNKNOWN | CY5 | NFQ-MGB | 26.516 |
| Non-scutellata E-2 | Target 2 | UNKNOWN | CY5 | NFQ-MGB | 26.571 |
| Non-scutellata E-2 | Target 2 | UNKNOWN | CY5 | NFQ-MGB | 27.121 |
| Non-scutellata E-3 | Target 2 | UNKNOWN | CY5 | NFQ-MGB | 30.362 |
| Non-scutellata E-3 | Target 2 | UNKNOWN | CY5 | NFQ-MGB | 30.402 |
| Non-scutellata E-3 | Target 2 | UNKNOWN | CY5 | NFQ-MGB | 30.723 |
| Non-scutellata E-4 | Target 2 | UNKNOWN | CY5 | NFQ-MGB | 34.839 |
| Non-scutellata E-4 | Target 2 | UNKNOWN | CY5 | NFQ-MGB | 35.383 |
| Non-scutellata E-4 | Target 2 | UNKNOWN | CY5 | NFQ-MGB | 36.322 |

**Supplementary detailed discussion of specific results**

***Cytb marker (qPCR assay II)***

*FABIS/USDA-ID bees*

We included four samples that had been previously identified as African derived honey bees (AHBs) using FABIS, and three of which were also identified as AHBs using USDA-ID. Sample ID: 121 was African/Africanized (FABIS-ID/USDA-ID), and our data supported this identification, with the sample being A-lineage and positive in our AHB test. Sample ID: 120 and 122 were both identified using FABIS as African and using USDA-ID as Introgression European (Table S1). Our data support this identity for Sample ID: 120, as we found it to be non-A-lineage. However, our data suggest that Sample ID: 122 is A-lineage and has the AHB mitotype for which we tested. Sample ID: 123 was identified using FABIS as AHB (no USDA-ID completed), and our data found it to be non-A-lineage. Sample ID: 124 served as a positive control consisting of honey bees from the HBREL apiary and assumed to be of European ancestry. This sample returned as non-A-lineage, suggesting European mitochondrial ancestry.

*Defensive samples from United States*

Of our defensive samples from Texas, one colony was determined to be non-A-lineage (Sample ID: 109, 133, 134) and the second colony was A-lineage, but not our AHB mitotype (Sample ID: 110, 125, 126). The sixteen defensive samples from Arizona came from five different colonies. One colony was determined to be non-A-lineage (Sample ID: 113, 130, 137, 431) and four were A-lineage. Of these four, only one colony showed the AHB mitotype (Sample ID: 132, 144, 145).

*South African samples*

The 18 South African samples that were positive for the AHB marker were from Bloemfontein, Citrusdal, Moorreesburg, Oudtshoorn, Springbok, Stellenbosch and Upington. The 19 *A. m. capensis* samples were from Bredasdorp, Cape Town, Citrusdal, East London, George, Oudtshoorn, Riversdale, Stellenbosch, Swellendam, Touwsrivier. These locations all fall within the known range of *A. m. capensis*, and the hybrid zone. For a map of these sampling locations, see (Eimanifar et al. 2018b). Twelve of the remaining 16 samples were A-lineage, but not AHB or *A. m. capensis*. Geographically, these came from locations across South Africa: Bredasdorp, East London, George, Pretoria, Stellenbosch, Swellendam, and Vryburg. The remaining four samples were also not AHB or *A. m. capensis*, but *qPCR assay I* was not run on these samples and so we cannot confirm their lineages (Table S1, assumed *A-lineage* based on RFLP ND2-PacI assay).
