## Supplementary figures and images for "Development of molecular markers for western honey bee (*Apis mellifera* L.) subspecies of regulatory concern in the United States"

### Figure S1

***Cytochrome b***

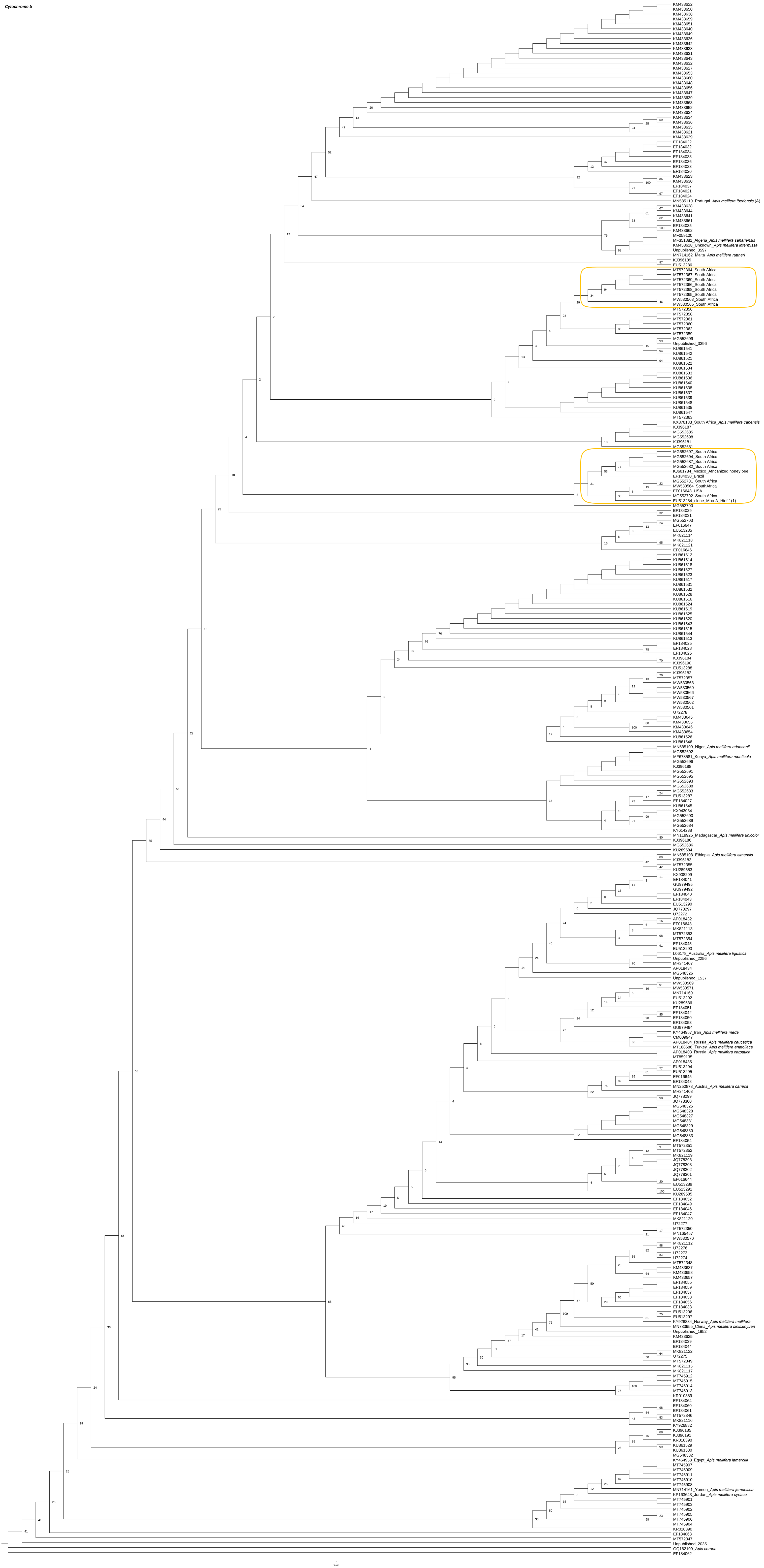

### Figure S2

**NADH dehydrogenase 2**

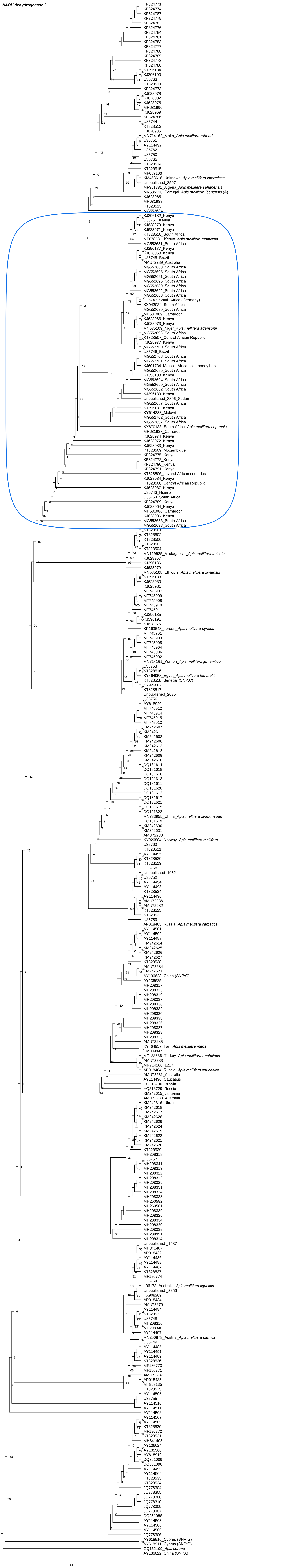
